# Shared safety abolishes the recovery of learned threat

**DOI:** 10.1101/2020.05.01.072827

**Authors:** Yafeng Pan, Andreas Olsson, Armita Golkar

**Affiliations:** Department of Clinical Neuroscience, Karolinska Institutet, Stockholm, Sweden; Department of Psychology, Stockholm University, Stockholm, Sweden

**Keywords:** social interaction, social learning, shared safety, recovery, threat

## Abstract

Social learning offers an efficient route to transmit information about threat and safety. To better isolate the processes that contribute to the efficacy of social safety learning, we developed a novel dyadic model of associative threat and extinction learning. In three separate social groups, we manipulated whether safety information during extinction was acquired via direct exposure to the conditioned stimulus (CS) in the presence of another individual (Direct exposure), via observation of other’s safety behavior (Vicarious exposure), or via the combination of both (Shared exposure). These groups were contrasted against a fourth group receiving direct CS exposure alone (Asocial exposure). Based on skin conductance responses, we observed that all social groups outperformed asocial learning in *inhibiting* the recovery of threat, but only Shared exposure *abolished* threat recovery. These results suggest that social safety learning is optimized by a combination of direct exposure and vicariously transmitted safety signals.

**Statement of relevance:** Humans, like other social animals, learn about threats and safety in the environment through social cues. Yet, the processes that contribute to the efficacy of social safety learning during threat transmission remain unknown. Here, we used a two-person approach to analyze skin conductance responses as participants engaged in a standard fear conditioning procedure (acquisition, extinction, and reinstatement). We found that during extinction, both (*i*) direct (conditioned stimulus) exposure in the presence of another individual and (*ii*) vicarious safety signals alone is sufficient to inhibit subsequent threat recovery, but that abolishing the recovery of conditioned threat responses requires a combination of both. This study has relevance for understanding how social information can optimize standard, asocial safety learning procedures to augment the effects of exposure on previously acquired fears. Thus, our work might help identify psychological and social strategies that can be used to counteract maladaptive fears in humans.

## 1. Introduction

Learning to predict and respond to threats and potential dangers in the environment is critical to adaption and survival. Such learning has been well described using associative threat learning models in both humans and animals (LeDoux, 2000), and has advanced our understanding of the etiology of threat and anxiety related disorder. These models are however inherently asocial as they rely on individual trial and error learning - neglecting that humans are inherently social. Most of us spend the majority of our daily lives socially connecting with others; these behaviors constitute social forms of learning, facilitating our acquisition of knowledge about the surrounding environment and influence our expectations and behaviors (Bandura, 1977; Olsson & Phelps, 2007).

Previous studies on social learning have mainly focused on how information about threats and potential dangers in the environment is transmitted between individuals (Debiec & Olsson, 2017; Lindström, Selbing, & Olsson, 2016) – a process known as *social threat learning* (LeDoux, 2017). Yet, less is known about how these learned threat responses can be changed and modified through socially acquired safety information. This is critical as deficits in the capacity to adapt and regulate responses to threatening cues when they no longer signal danger is a key feature of threat and anxiety-related disorders (Graham & Milad, 2011). Support for the utility of exploiting social information to regulate aversive responses mainly comes from research demonstrating that the presence of a conspecific attenuates the behavioral, physiological, and neural response to a stressful or threat-provoking event. This “social buffering” phenomenon has been found across social species, including zebrafish (Faustino, Tacão-Monteiro, & Oliveira, 2017), rodents (Kiyokawa, Takeuchi, & Mori, 2007), non-human primates (Wittig et al., 2016), and humans (Ditzen & Heinrichs, 2014; Gunnar & Hostinar, 2015), suggesting the existence of an evolved system by social interaction can modulate behavior.

More specifically, previous experimental work in humans have demonstrated that acute physical (Roberts, Klatzkin, & Mechlin, 2015) and social stress (Heinrichs, Baumgartner, Kirschbaum, & Ehlert, 2003) result in significantly smaller elevations in heart rate, blood pressure, and cortisol in humans socially supported by another partner, compared to individuals who undergo stress alone. More recently, the mere presence of another partner was shown to reduce psychophysiological responses to aversive sounds, compared to an alone condition (Qi et al., 2020). The social buffering research has however almost exclusively focused on the social regulation of stress responses (Hostinar, Sullivan, & Gunnar, 2014) and little work has focused on how social information modulates learned threat memories or how social safety information is learned (but see also Ferreira et al., 2019). Briefly, extinction learning involves the formation of novel, safety associations after repeated safe exposures to a threatening stimulus; i.e. learning that the stimulus that earlier predicted danger now predicts safety. However, unlike threat learning, the threat-reducing effects of extinction learning decline with the lapse of time, do not generalize across different contexts, and are easily reinstated in behavior following re-exposure to cues associated with the original fear memory (Bouton, 2002). This recovery of threat responses after extinction also contributes to relapse of fear that is common after extinction-based exposure therapies that are used in the treatment of threat and anxiety-related disorders (Rachman, 1977). Consequently, one major aim of the research on threat learning and regulation is to use experimental extinction as a translational model to identify strategies that can prevent the recovery of threat after extinction in order to guide novel treatment approaches (Bouton, Mineka, & Barlow, 2001).

To address whether social information can optimize the effects of extinction learning and prevent the recovery of threat in humans, previous studies on both social threat and safety learning have commonly adopted video-based social learning paradigms (Golkar, Haaker, Selbing, & Olsson, 2016; Golkar, Selbing, Flygare, Ohman, & Olsson, 2013; Haaker, Golkar, Selbing, & Olsson, 2017). In these paradigms, the observer learns about threat or safety by watching a pre-recorded video depicting the behavior of a demonstrator reacting either fearfully or calmly in the presence of threat. Although the recent decade has seen a paradigm shift towards real-time interactions in both social psychological and neuroscientific research (Pan et al., 2020; Redcay & Schilbach, 2019), few studies in humans have addresses whether social learning of threat and safety generalizes to a more ecologically valid setting. This methodological limitation critically also constrains the possibility to isolate core social aspects of the learning situation that can unveil the processes that contribute to safety transmission between individuals.

In the current study, we developed a dyadic experimental model with the aim to isolate the social processes that contribute to safety transmission. Given that extinction learning serves as a translational model for threat and anxiety disorders, we focused particularly on how social safety learning can prevent the recovery of learned threat responses. Live pairs of participants underwent an associative threat learning and extinction procedure followed by a threat recovery test. To isolate the social processes that are necessary for the efficacy of social safety learning, we ran three separate social safety learning groups: we manipulated whether safety information was acquired via own conditioned stimulus (CS) exposure in the mere presence of another individual (*Direct exposure*), via observation of another individual’s safety behavior (*Vicarious exposure*) or via the combination of both (*Shared exposure*). These social safety learning groups were contrasted with a standard, asocial safety learning group who received direct CS exposure alone (*Asocial exposure*). Learning was indexed using skin conductance responses (SCRs) throughout all stages.

Our aim was to test two main predictions. First, to the extent that social buffering theory (Ditzen & Heinrichs, 2014; Gunnar & Hostinar, 2015) holds true, we would predict social safety learning (Direct, Vicarious, or Shared exposure) to outperform asocial safety learning in *inhibiting* the recovery of threat. Second, in accordance with previous literature on safety learning (Golkar & Olsson, 2016; Golkar, Selbing, et al., 2013), we predicted that Shared exposure, involving both CS and model exposure, would *abolish* threat recovery. For this second hypothesis, we restricted our analysis to the early trials of reinstatement because previous research showing that reinstated threat responses typically undergo rapid re-extinction (Haaker, Golkar, Hermans, & Lonsdorf, 2014). Finally, based on previous research showing that inter-personal physiological synchrony is associated with social behaviors (Palumbo et al., 2017), including threat learning (Pärnamets, Espinosa, & Olsson, 2020), we carried out an exploratory analysis to assess the physiological synchrony between two partners in a dyad during social safety learning.

## 2. Methods

### 2.1. Ethics statement

This study was conducted in accordance with guidelines and regulations laid down in the Declaration of Helsinki. Written informed consent was obtained from each participant prior to the experiment. The study procedure was approved by Regional Ethical Review Board in Stockholm (DNR: 2016/249-31/1). Each participant was reimbursed with two movie tickets for their participation.

### 2.2. Participants

Power calculations were performed based on the obtained effect size (d = 0.75) in our original study comparing social and asocial extinction learning (Golkar, Selbing, et al., 2013). Setting the threshold to 95% sensitivity (1-beta = 0.95) and a significance level of alpha = 0.05 yielded a required sample size of 40 in each group. In an endeavor to increase statistical power, we decided to oversample to reach a minimum of 50 participants in each experimental group. To achieve this, 231 participants were recruited from the student population at Karolinska Institutet. Participants were screened from having previously partaken in conditioning experiments and we confirmed that the participants in each dyad did not already know each other prior to participating in the experiment. Of the 231 participants who completed the experiment, we excluded 1 participant who failed to show any measurable skin conductance response (< 0.02 microsiemens, μS) to any CS during the entire experiment. This left 230 participants in the final sample. These participants were randomly assigned to either one of the three different social learning groups (Direct exposure, Vicarious exposure, and Shared exposure) or the asocial learning control group (for group characteristics and demographic information, see **Table1**).

**Table 1.**
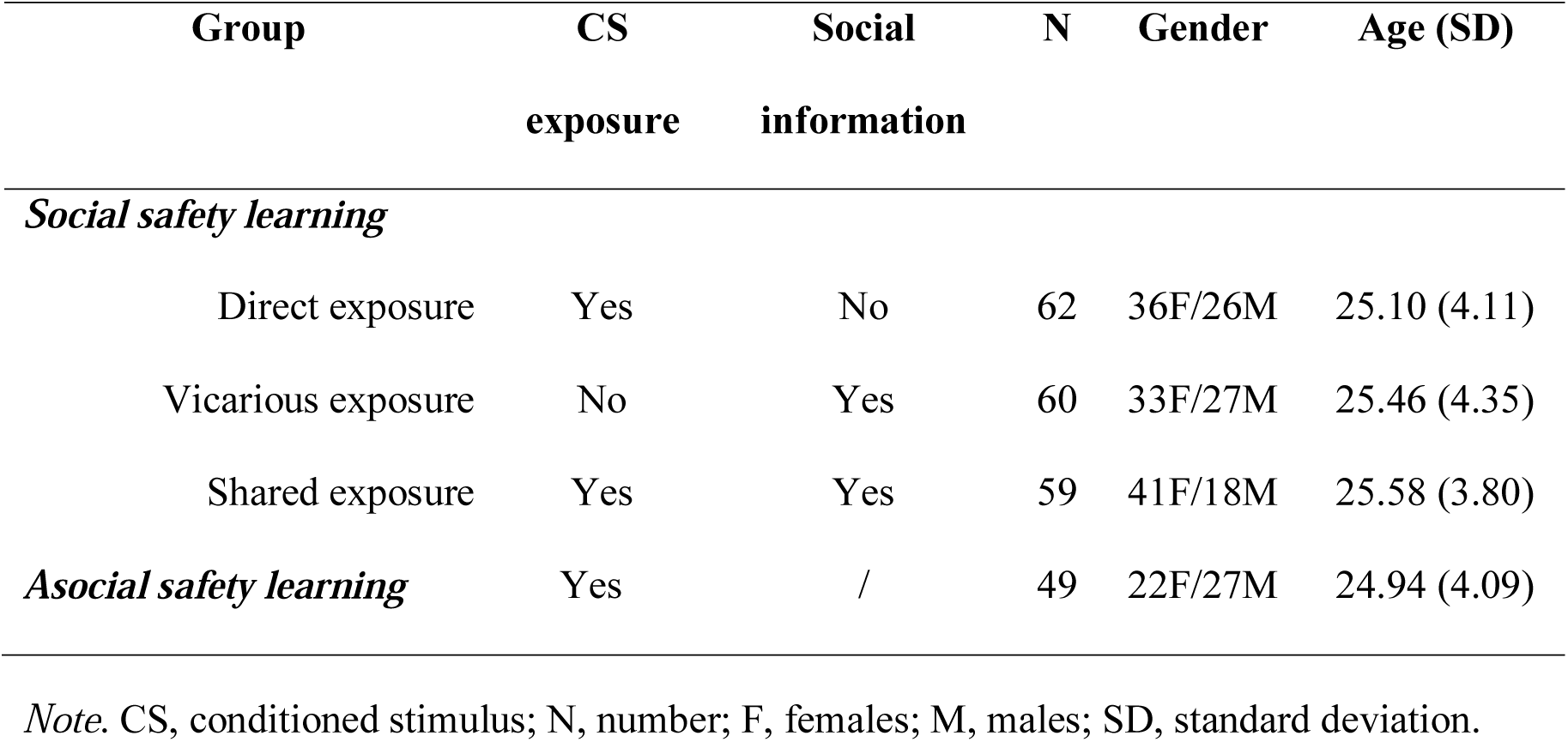
Group characteristics and demographic information.

### 2.3. Experimental set-up

#### Social safety learning

Participants were enrolled in pairs of two participants that did not report previous contact or familiarity with each other. Each pair were seated in a waiting area during and were given time to read and sign informed consent forms before participation.

Then, each participant underwent a standardized shock calibration alone, with the aim to individually adjust the level of shock intensity to a level that was highly uncomfortable but not painful. Next, they were seated next to each other in the experimental room separated by a moveable wall that occluded any visual information of the other participant. In addition, the monitors were equipped with privacy screen filters to ensure that the visual presentations were not visible to the other individual in each pair. Participants were also instructed they were not allowed to talk to each other during any parts of the experiment. They were also informed that there would be brief interruptions during which the experimenter would make some adjustments and that should refrain from asking questions during these breaks.

Prior to the experiment (see **Figure 1** for an overview of the experimental stages), all participants were allowed to familiarize with the stimuli. Participants were exposed to two non-reinforced presentations of two different stimuli (one spider image and one snake image) that would subsequently serve as the conditioned stimuli (CSs). The formal experiment consisted of the following three stages: acquisition, extinction, and reinstatement test.

**Figure 1.**
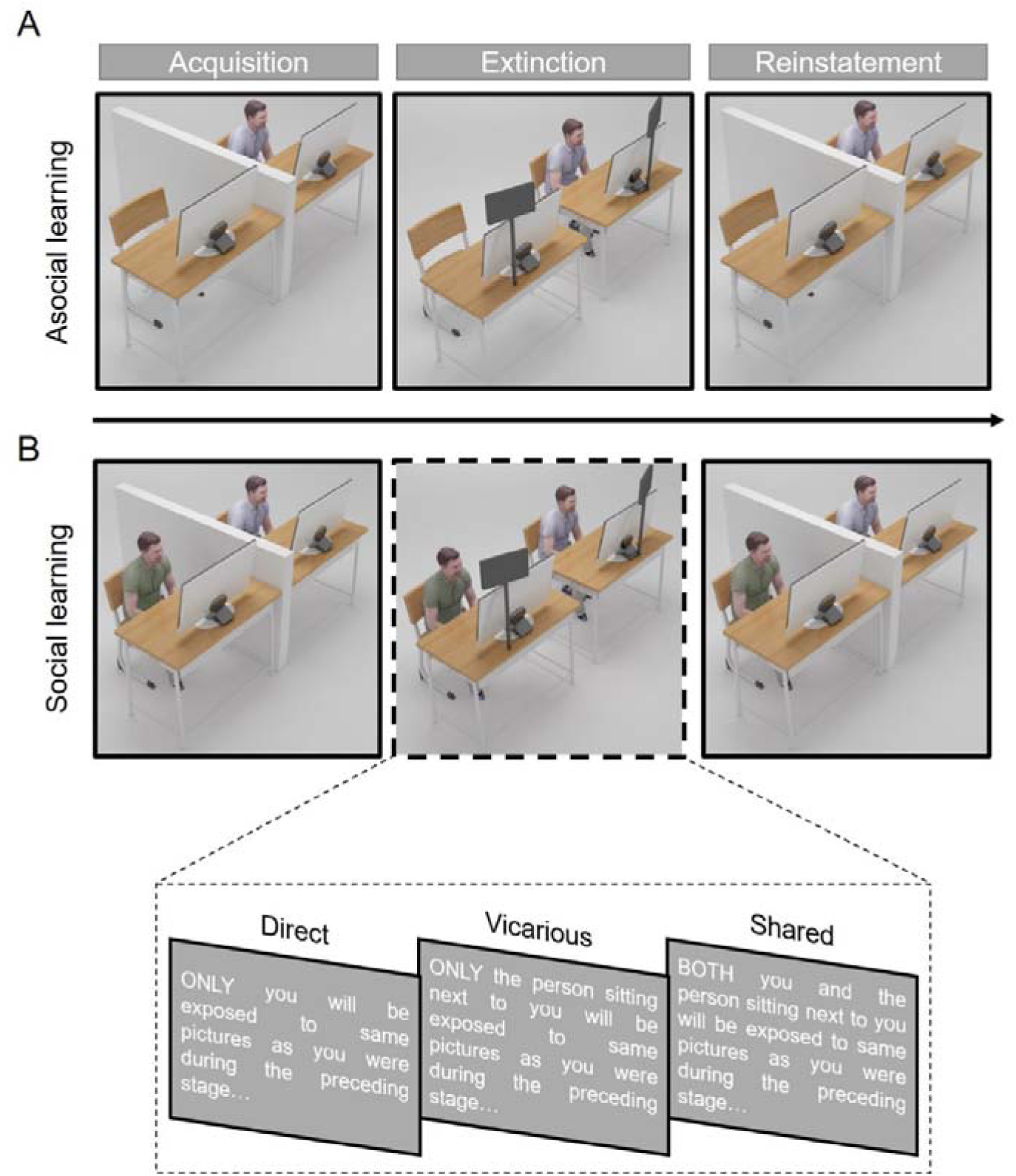
General set-up. All participants in (A) and (B) underwent an identical acquisition and reinstatement stage with the exception that in (A), the Asocial learning group performed all stages alone. (B) Participants in the social learning group were further categorized into three sub-groups (Direct, Vicarious, and Shared exposure) based on different CS exposure and instructions they received during extinction. Participants in both the Direct exposure and Shared exposure groups viewed the same CS as during the previous stage but either instructed that the person sitting next to them were observing novel, irrelevant stimuli (Direct exposure) or the same stimuli (Shared exposure) as themselves. Participants in the Vicarious exposure group were exposed to two novel, neutral stimuli and instructed that only the person sitting next to them were exposed to the stimuli from the preceding stage. During all stages, screen filters occluding all information of what the other participant was exposed to were used.

During the acquisition stage, all participants underwent a standard threat acquisition protocol involving two CSs. For each participant, one CS predicted the delivery of a shock (the unconditioned stimulus; US) on 2/3 of the CS presentations, whereas the other CS (CS-) never predicted the US. This acquisition stage ended with an instruction screen: “The experimenter will not make some minor adjustment, please remain seated”.

During the subsequent extinction stage, the experimenter removed the movable wall between the participants and adjusted mirrors that allowed the participants to receive visual feedback from the face of the other participant. The two mirrors were placed in an angle so that the reflection of the other participant’s face did not require them to move their gaze away from the visual presentation on their screen. We manipulated safety transmission in three separate groups by changing both CS exposure and instructions. In the *Direct exposure* group, participants received the following written instructions: “During the next stage, ONLY you will be exposed to same pictures as you saw during the preceding stage, and the person sitting next to you will be exposed to two novel pictures. In the *Vicarious exposure* group, participants were informed that “During the next stage, you will be exposed to a presentation of two novel pictures, and ONLY the person sitting next to you will be exposed to the same pictures as you were exposed to during the preceding stage” and the *Shared exposure* group were informed “During the next stage, BOTH you and the person sitting next to you will be exposed to the same pictures as during the preceding stage”. For the Direct and Shared exposure groups, the extinction stage consisted of non-reinforced presentations of each CS. In the Vicarious exposure group, the CSs were replaced by the presentation of two neutral and non-related control stimuli (picture of a butterfly and a bird).

During the final reinstatement stage, the experimenter re-positioned the wall between the participants and removed the mirrors. After this readjustment, all participant received identical instruction on their screen “The experiment will now be resumed” which was followed by a standard *reinstatement* test during which participants received 3 un-signaled shocks followed by a black screen (34 s) before all CSs were represented to the participants.

#### Asocial safety learning

Participants underwent the same experimental stages (acquisition, extinction, and reinstatement) using the same set-up but alone in the absence of another participant. After the acquisition stage, participants in this *Asocial exposure* group received the following instructions “During the next stage, you will now be exposed to the same two pictures as you were during the preceding stage”. This stage was followed by the reinstatement stage.

During all stages of the experiment, each CS was presented six times with a duration of 6 s. Between each CS trial, a black screen (i.e., inter-stimulus interval, ITI) was presented for 12–18 s.

#### Manipulation check

All pairs reported seeing the other participants face in the mirrors during extinction, without having to divert gaze from the CS presentation, and none of the included participants reported being able to see the CS presentation of the other individual.

### 2.4. Data acquisition

Participants’ SCRs were recorded and measured using a Biopac MP150 system and AcqKnowledge 4.0 software (BIOPAC Systems Inc., Goleta, CA, USA). Ag-AgCl electrodes were attached to the middle and index fingers from the hand not receiving shocks. A conducting gel was used to improve signal quality, in accordance with standard recommendations (Boucsein, 2012). Electric shocks (aversive stimuli) were delivered using a STM200 module (BIOPAC Systems Inc., Goleta, CA, USA). The sampling rate was set to 2 kHz. Shocks consisted of a single 100ms DC pulse, which were applied to the lower forearm. The intensity of the shocks was individually calibrated, following the rule being “unpleasant but not painful”. Stimulus presentation and shock delivery were controlled by the Presentation software (Neurobehavioral Systems, Inc., Albany, CA).

### 2.5. Data analysis

SCRs were continuously recorded throughout all experimental stages (acquisition, extinction, and reinstatement test). The raw signals were filtered offline with a low-pass filter (1 Hz) and a high-pass filter (0.05 Hz). The SCR was measured for each stimulus trial as the base-to-peak amplitude of the first response (in μS) in the latency window from 0.5 to 4.5 s following stimulus onset (Haaker et al., 2017). Minimal response criterion was set to 0.02 μS, and responses that did not pass this criterion were scored as 0. SCR scores were then *z*-transformed to normalize the distributions (Boucsein, 2012). Clean SCRs were then used for subsequent statistical comparisons.

In addition, to access the physiological synchrony between two participants in a dyad, clean SCR time series was further submitted to an exploratory analysis using the Pearson correlation coefficient (Piazza, Hasenfratz, Hasson, & Lew-Williams, 2020; Reddan, Young, Falkner, López-Solà, & Wager, 2020). The correlation coefficients between two SCRs from participant # 1 and participant #2 were calculated over the whole extinction stage (**Figure 1**). Resulting values were Fisher-z-transformed. Note that only Social exposure groups (Direct, Vicarious, and Shared exposure) were included in this exploratory analysis, as physiological synchrony should be estimated within dyads. One dyad containing a missing participant (failed in SCR acquisition, see section 2.2) were excluded, resulting in 31 pairs in the Direct exposure group, 30 pairs in the Vicarious exposure group, and 29 pairs in the Shared exposure group.

### 2.6. Statistical analyses

All the statistical analyses were conducted using IBM SPSS (version 21, SPSS Inc., Chicago, IL, USA). Each stage of the experiment was analyzed separately and significant interactions were followed up by separate two-tailed *t*-tests. Significance level was set to *p* < 0.05. η^*2*^ and Cohen’s *d* are reported as measures of effect size where appropriate.

#### 2.6.1. Confirmatory analysis

When analyzing data from the acquisition stage, a 2 × 6 × 4 mixed-design Analysis of Variance (ANOVA), with Stimulus (CS+ and CS-) and Trial (#1∼6) as two within-subject factors and Group (Direct, Vicarious, Shared, and Asocial exposure) as a between-subject factor, was used to analyze all results across conditions. Degrees of freedom were corrected using Greenhouse-Geisser estimates when assumption of sphericity was violated. We predicted successful threat acquisition in all experimental groups. Successful acquisition was defined as higher SCR to the CS+ relative to the CS-across acquisition, which would be supported by a significant Stimulus × Trial interaction in the absence of any significant effects involving groups.

Data from the extinction stage were analyzed using the same mixed-design ANOVA. Extinction of the conditioned SCR was defined as decreasing SCR responses to the CS+ relative to the CS-across extinction training (Stimulus × Trial interaction). We anticipated a decreasing trend of CS+/CS- differentiation (as a marker of diminishing conditioned SCR) across trials in all groups except in the Vicarious exposure group that did not receive CS exposure during extinction and was therefore not expected to show significant CS+/CS- differentiation during this stage (no CS exposure, see **Table 1**).

#### 2.6.2. Hypothesis-driven analysis

When analyzing data from the reinstatement stage, we carried out a series of statistics in a hypothesis-driven way. To test our first prediction i.e., that social safety learning would produce less recovery of learned threat compared to asocial safety learning, we compared CS+/CS- differentiation across reinstatement in a Stimulus (2) × Trial (6) × Group (4) mixed-design ANOVA, followed by planned comparisons between each social exposure group (Shared, Vicarious, or Direct) with the Asocial exposure group. To test our second hypothesis (i.e., that the Shared exposure group would abolish CS+/CS- differentiation during early reinstatement), we ran a Stimulus (2) × Trial (2: first 2 trials) × Group (4) mixed-design on SCR. We only focused on the early trials of reinstatement due to rapid re-extinction during reinstatement (Haaker et al., 2014), and in line with previous research demonstrating the efficacy of social safety learning in preventing the recovery of threat responses focusing on these early trials (Golkar, Castro, & Olsson, 2015; Golkar et al., 2016; Golkar, Tjaden, & Kindt, 2017). Each set of planned contrasts were Bonferroni corrected to account for multiple comparisons.

#### 2.6.3. Exploratory analysis

Finally, we conducted exploratory analyses to test the potential physiological synchrony between two participants in a dyad under social settings (i.e. during extinction). Synchrony was estimated using the Pearson correlational approach, in accordance with recent advances (Piazza et al., 2020; Reddan et al., 2020). To evaluate the statistical significance of the Pearson correlation coefficients at the extinction stage, we created a null distribution based on a surrogate dataset. The surrogate dataset was obtained by Fourier transforming the original SCR signal, randomizing the phases of Fourier coefficients, performing an inverse Fourier transform, and re-computing the correlation coefficients (Liu et al., 2017). This procedure was repeated 1,000 times. Significance levels (thresholded at 0.05) were evaluated by comparing the correlation coefficients from the original dataset and those from the surrogate dataset. It was assumed that physiological synchrony based on original dataset should be higher than that based on surrogate dataset, as the latter disrupts the temporal structure of the original signals.

## 3. Results

### 3.1. Confirmatory results

#### 3.1.1. Acquisition stage

First we confirmed that all group successfully acquired conditioned threat responses as supported by significantly higher SCR to the CS+ than the CS-across acquisition (Stimulus: *F*_1,226_ = 13.74, *p* < 0.001, η_*partial*_^2^ = 0.06; Stimulus × Trial interaction: *F*_4.71,1063.55_ = 37.08, *p* < 0.001, η_*partial*_^2^ = 0.14; see **Figure 2** (left column). There were no significant effects involving Group (main effect: *F*_3, 226_ = 0.14, *p* = 0.94; Trial × Group: *F*_12.93,974.11_ = 1.17, *p* = 0.30; Stimulus × Group: *F*_3, 226_ = 2.15, *p* = 0.10; Stimulus × Trial × Group, *F*_14.12,1063.55_ = 0.40, *p* = 0.98).

**Figure 2.**
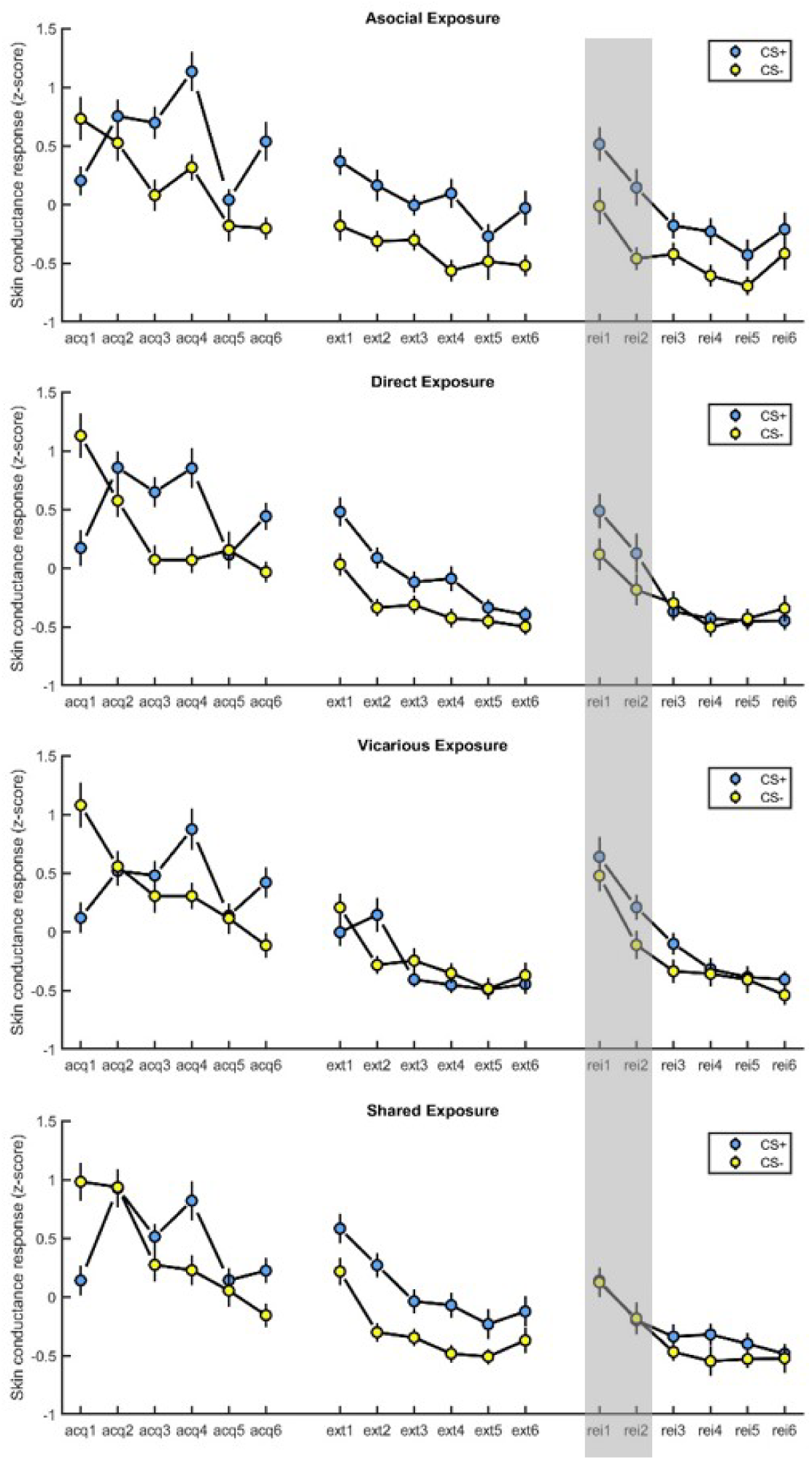
Mean skin conductance responses (z-score) across stimuli, groups, and trials over the entire experiment. The graphs from upper to bottom represent the data from Asocial exposure, Direct exposure, Vicarious exposure, and Shared exposure groups, respectively. Left column: acq1-acq6 refers to the 6 acquisition trials; middle column: ext1-ext6 refers to the 6 extinction trials; right column: rei1-rei6 refers to the 6 reinstatement trials. Grey shadow highlights the findings from the early reinstatement stage, indicating that shared safety abolishes the recovery of learned threat. Error bars denote standard errors.

#### 3.1.2. Extinction stage

Second, we confirmed that CS+/CS- differentiation decreased across extinction. Progressive extinction was confirmed by a main effect of Trial (*F*_4.19,946.85_ = 47.66, *p* < 0.001, η _*partial*_^2^ = 0.17) and an interaction of Stimulus × Trial (*F*_4.58,1034.82_ = 3.49, *p* = 0.01, η _*partial*_^2^ = 0.02), with a generally decreasing trend in CS+/CS- differentiation (**Figure 2**, middle column). However, as expected, CS+/CS- differentiation differed between the Vicarious exposure group that did not receive own CS exposure and the other groups. This was evidence by a main effect of Group (*F*_3,226_ = 3.87, *p* = 0.01, η_*partial*_^2^ = 0.05) and a Stimulus × Group interaction (*F*_3,226_ = 9.14, *p* < 0.001, η _*partial*_^2^ = 0.11), indicating that CS+/CS- differentiation was significant in all groups (Asocial exposure, *t*_48_ = 4.66, *p* < 0.001, Cohen’s *d* = 2.14; Direct exposure, *t*_61_ = 4.61. *p* < 0.001, Cohen’s *d* = 0.85; and Shared exposure, *t*_58_ = 6.39, *p* < 0.001, Cohen’s *d* = 1.03), except for the Vicarious exposure group that did not receive own CS exposure (*t*_59_ = 0.42, *p* = 0.68). This analysis did not yield a significant three-way interaction (*F*_13.74,1034.82_ = 1.06, *p* = 0.39).

To further elucidate the extinction effect, we analyzed CS+/CS- differentiation over the last two trial of the extinction stage. Results indicated that the Direct exposure (*F*_1,61_ = 1.92, *p* = 0.17) and Vicarious exposure (*F*_1,59_ = 0.27, *p* = 0.61) groups efficiently extinguished CS+/CS- differentiation, whereas the Shared exposure (*F*_1,58_ = 6.12, *p* = 0.02, η_*partial*_^2^ = 0.10) and Asocial exposure (*F*_1,48_ = 5.40, *p* = 0.02, η_*partial*_^2^ = 0.10) groups did not.

### 3.2. Hypothesis-driven results

To test our first prediction that all social safety learning groups would better inhibit reinstatement (i.e., less CS+/CS- differentiation) compared to the Asocial exposure group, we ran a Stimulus (2) × Trial (6) × Group (4) mixed design ANOVA. Overall, this analysis yielded a significant Stimulus × Group interaction (*F*_3,226_ = 3.40, *p* = 0.02, η_*partial*_^2^ = 0.04), a main effect of Group (*F*_3,226_ = 3.22, *p* = 0.02, η _*partial*_^2^ = 0.04) and no interaction with Trial (*F*s < 1.03, *p*s > 0.42). Comparisons of the separate social exposure groups (Shared, Vicarious, or Direct) vs. the Asocial exposure group using adjusted alpha levels of *p* = 0.017 (0.05/3) resulted in a significant Stimulus × Group interaction in the Shared vs. Asocial exposure comparison (*F*_1,106_ = 7.64, *p* = 0.01, η _*partial*_^2^ = 0.07), which, as predicted, was driven by lower CS+/CS- differentiation in Shared exposure relative to Asocial exposure (see **Figure 2**, right column). Corresponding analyses for the Vicarious vs. Asocial exposure group (*F*_1,107_ = 5.08, *p* = 0.01, η_*partial*_^2^ = 0.05) and the Direct vs. Asocial exposure (*F*_1,109_ = 7.38, *p* = 0.01, η_*partial*_^2^ = 0.06) were also significant at the adjusted alpha level.

To test our second prediction that the Shared exposure would abolish CS+/CS- differentiation during early reinstatement, we next analyzed early reinstatement test performance. This analysis resulted in a significant main effect of Group (*F*_3,226_ = 3.64, *p* = 0.01, η_*partial*_^2^ = 0.05), and Trial (*F*_1,226_ = 31.00, *p* < 0.001, η _*partial*_^2^ = 0.12) and a significant Stimulus × Group interaction (*F*_3,226_ = 2.86, *p* = 0.036, η_*partial*_^2^ = 0.04). Using adjusted alpha levels of *p* = 0.017 (0.05/3), we observed the predicted difference between the Shared exposure and the Asocial exposure group (Stimulus × Group, *F*_1,106_ = 9.19, *p* = 0.003, η_*partial*_^2^ = 0.08), and no significant differences between Vicarious vs. Asocial exposure (*F*_1,107_ = 3.55, *p* = 0.06), or Direct vs. Asocial exposure (*F*_1,109_ = 1.32, *p* = 0.25). Critically, in contrast to Asocial exposure (main effect of Stimulus, *F*_1,48_= 23.21, *p* < 0.001, η _*partial*_^2^ = 0.33), Shared exposure abolished CS+/CS- differentiation during early reinstatement (main effect of Stimulus, *F*_1,58_ = 0.00, *p* = 0.99, **Figure 3A**) using an adjusted alpha level of *p* = 0.025 (0.05/2). The absence of CS+/CS- differentiation during early reinstatement in the Shared Exposure group was further evidenced by a Bayesian paired samples *t*-test using JASP software (https://jasp-stats.org/, JASP Team, 2017), 95% Credible Interval = [-0.30, 0.21], Bayes Factor (BF_10_) = 0.16 (**Figure 3B**).

**Figure 3.**
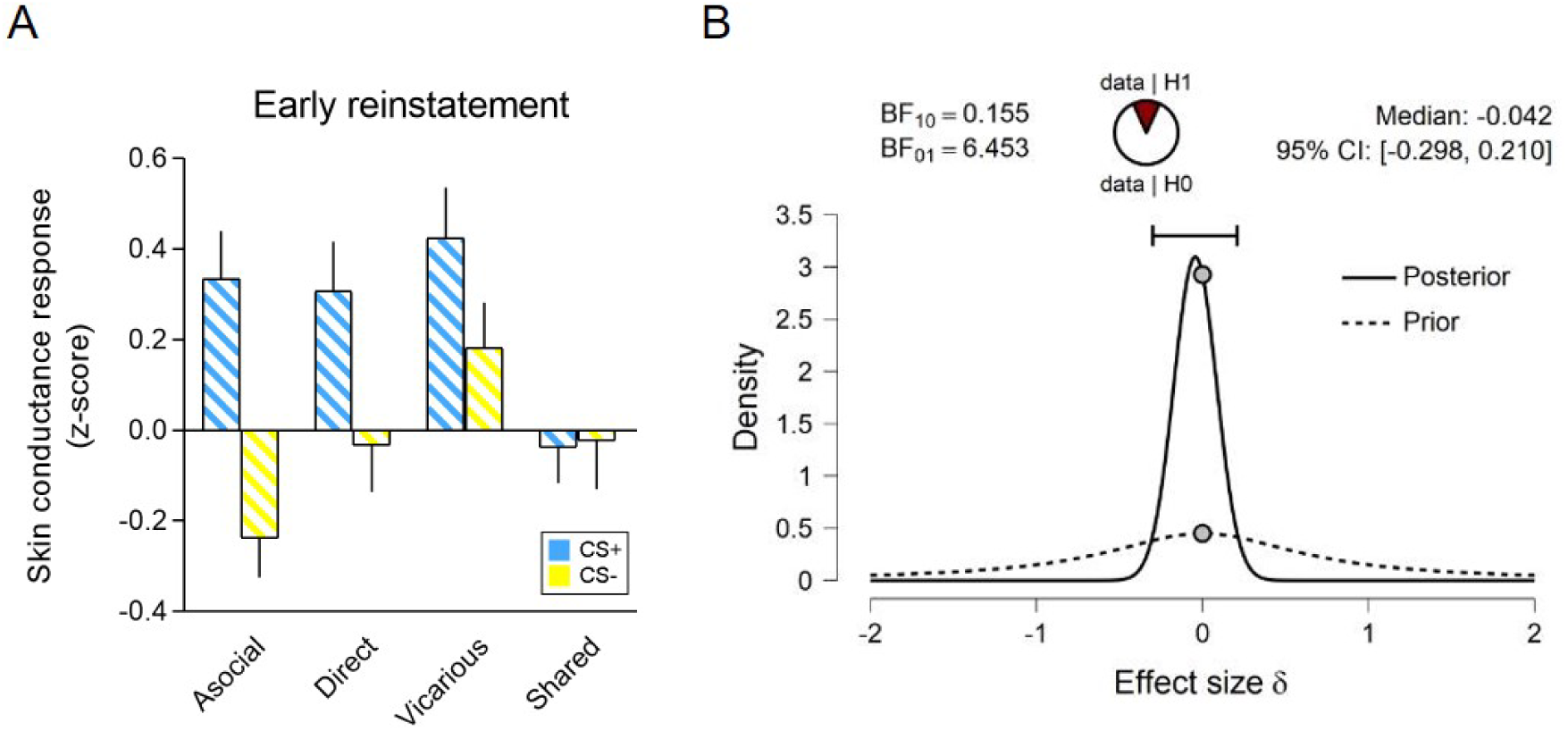
(A) Mean skin conductance response (z-score) across stimuli, groups during the early reinstatement stage. Error bars represent standard errors. (B) Inferential plots for prior and posterior probabilities. Skin conductance responses to CS+ and CS- during early reinstatement in the Shared Exposure group were compared using a Bayesian paired samples *t*-test.

### 3.3. Exploratory results

Our exploratory analysis focused on evaluating the physiological synchrony between two participants in a dyad at the extinction stage (**Figure 4**), during which the dyads were situated in a real-time social setting. All three social safety learning groups showed significantly stronger synchrony (averaged across trials and stimuli) between the SCR signals of two participants within dyads compared to a surrogate dataset (generated by phase-scrambling of original signals): Direct exposure (mean ± SD, 0.19 ± 0.05), Vicarious exposure (0.08 ± 0.05), and Shared exposure (0.20 ± 0.05), *p*s < 0.02.

**Figure 4.**
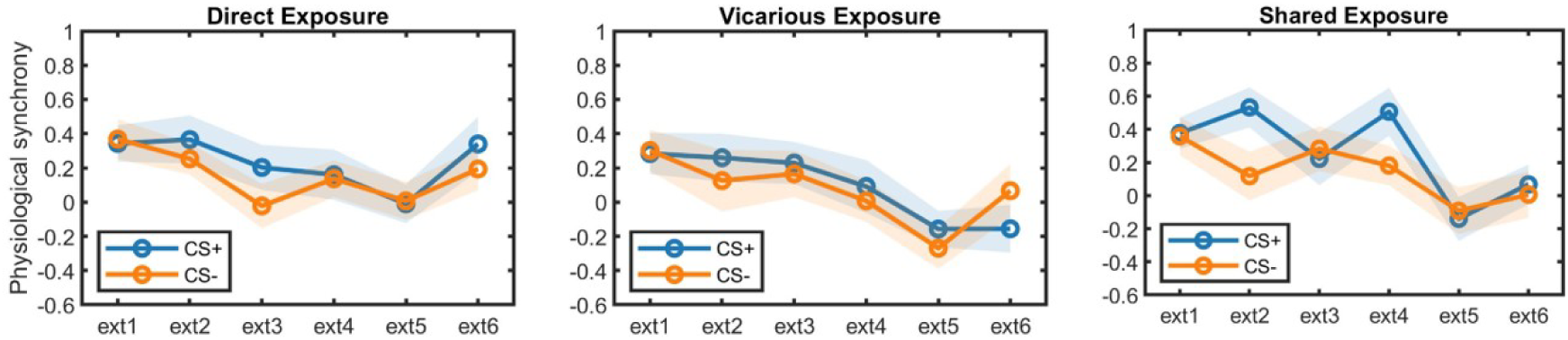
Physiological synchrony (average *r*) between skin conductance responses from two participants in a dyad across groups during the extinction stage. Ext1-ext6 refers to the 6 extinction trials. Shadow areas represent ± 1 standard errors.

An additional Stimulus (2) × Trial (6) × Group (3) mixed-design ANOVA on physiological synchrony revealed a main effect of Trial, *F*_4.46,388.12_ = 7.66, *p* < 0.001, η_*partial*_^2^ = 0.08, suggesting that synchrony generally attenuated across trials. The main effect of Stimulus was marginally significant, *F*_1,87_= 3.11, *p* = 0.08, η _*partial*_^2^ = 0.04, indicating that synchrony was more pronounced for CS+ vs. CS-. No other significant effects were observed (*F*s < 1.91, *p*s > 0.15).

## 4. Discussion

This study investigated how the recovery of learned threat responses are influenced by socially acquired safety information. In doing so, the study departed from two main hypotheses: (*i*) that social safety learning outperforms asocial safety learning in *inhibiting* the recovery of learned threat responses, and (*ii*) shared social safety learning *abolishes* threat recovery during early reinstatement. To address these hypotheses, we used a standard conditioning procedure (Golkar, Selbing, et al., 2013), which consists of three stages: (*i*) acquisition –threat acquisition through direct CS-US exposure; (*ii*) extinction –safety learning in the absence of aversive outcome (no US); and (*iii*) reinstatement – recovery of learned threat to the CSs after re-exposure to the US. Our results show that (*i*) compared to asocial safety learning, all three social safety learning groups were more successful in inhibiting the recovery of learned threat responses, as measured during a subsequent reinstatement stage, and (*ii*) only shared safety learning abolished the recovery of learned threat response, as evidenced by the non-significant CS+/CS- differentiation (confirmed by Bayesian factor analysis) during early reinstatement. These results suggest that both direct CS exposure in the presence of another person, and vicarious safety signals alone is sufficient to produce better inhibition of the recovery of learned threat responses compared to asocial learning, but that during early reinstatement, the beneficial effects of socially acquired safety relative to asocial safety learning are particularly pronounced when safety signals can be acquired from both direct experience and from another person (*Shared exposure*).

Previous studies supporting a modulatory role of social signals on aversive responses come mainly from non-human animal work (Cohen & Wills, 1985) and specifically from rodent models that have demonstrated that the expression of conditioned threat responses are reduced in the presence of a non-fearful conspecific (Ferreira et al., 2019). This previous work indicates that a key factor facilitating safety learning is a social modulation mechanism, which requires the presence of a social conspecific and the trigger of safety signals during safety learning. Such “social modulation” might be further empirically characterized in terms of physiological synchrony (Palumbo et al., 2017). We observed higher synchrony for all social learning groups, compared to phase-scrambled controls. Thus, physiological synchrony might support shared experience and/or interpersonal information transfer by aligning physiological processes between the participants (Pärnamets et al., 2020). This study did not find any significant difference in physiological synchrony between the social groups; the lack of relationship deserves further investigations, as it may reflect either a lack of sensitivity regarding paradigms or analogous effects of social modulation on interpersonal autonomic coupling during safety learning.

The interpretation of our findings requires some considerations. First, the augmented inhibition of threat recovery during early reinstatement in the Shared exposure group does not seem to be related to end extinction performance. Neither Shared exposure nor Asocial exposure entirely abolished CS+/CS- differentiation during end extinction, but strikingly, they differed during the subsequent reinstatement stage. This lack of consistency between end extinction performance and recovery phenomena has been documented in previous research (Golkar, Bellander, & öhman, 2013; Prenoveau, Craske, Liao, & Ornitz, 2013). Second, an inspection of **Figure 3A** shows that whereas both the Shared and Vicarious extinction group showed relatively low CS+/CS- differentiation during early reinstatement, the lack of CS+/CS- differentiation in the Shared exposure group was due to comparably lower CS+ responses. In contrast, the Vicarious exposure group showed high CS+ *and* high CS- responses, indicating generalization of conditioned threat responses in this group. The stronger CS- response observed in the Vicarious exposure group might be related to the fact that this group did not receive CS exposure during the extinction stage and therefore showed overall stronger SCRs to the re-appearance of the CSs. Third, the current study did not allow for verbal interaction between two individuals. However, to ensure that participants could access socially transmitted information, we set-up mirrors so participants could observe each other’s emotional reactions (e.g., facial expression and body language) during the safety learning (i.e., extinction) stage. Fourth, the results from the exploratory analyses should be interpreted with caution, as our power calculations were based on a priori hypotheses for the main analyses of threat-related SCR responses. An inevitable consequence for the analyses focusing on physiological synchrony between dyads is that the sample size is reduced by 50% - which represents a common issue in dyadic studies (Pan & Cheng, 2020).

Taken together, these observations indicate that both direct CS exposure in the physical presence of another person, and the mere safety signal from another person alone is sufficient to inhibit the recovery of learned threat responses. Abolishing the recovery of threat, however, seems to require the combination of both (or beyond). As we did not have an a priori hypothesis for differences between the social safety learning groups (Direct vs. Vicarious vs. Shared exposure; i.e., which group best inhibits the recovery of learned threat), this question should be further addressed in future research powered enough to detect between-group differences. Nevertheless, to our knowledge, our work represents the first demonstration of social safety learning in a two-person situation with high ecological validity. The current set of findings extends recent research addressing the influence of physical presence on physiological responding to aversive stimuli (Qi et al., 2020) by addressing the influence of social modulation of threat in a safety learning context. More specifically, by focusing on safety learning, this work has relevance for understanding how social information can optimize standard, asocial safety learning procedure to augment the effects of exposure on previously acquired fears. Future research should more closely address the relative contribution of different social processes to safety learning and delineate the underlying mechanistic features that characterize these processes.

## Data accessibility

The full dataset used is accessible at https://osf.io/h3ckv/.

## Authors’ contributions

Armita Golkar conceived the study and designed the experiment with input from Andreas Olsson. Armita Golkar and Yafeng Pan performed the statistical analysis and interpreted the data and Yafeng Pan drafted the manuscript. All authors critically revised the manuscript and gave final approval for publication and agree to be held accountable for the work performed therein.

## Acknowledgements

We would like to thank Lisa Espinosa, Cathelijn Tjaden and Anastasia Dima for help with the data collection and scoring. This research was supported by a Consolidator Grant (2018-00877) from the Swedish Research Foundation (Vetenskapsrådet), the Knut and Alice Wallenberg Foundation (KAW 2014.0237), and a Starting Grant (284366) from the European Research Council to Andreas Olsson, and an independent project grant to Armita Golkar from the Swedish Research Council (Vetenskapsrådet: 2017-02210).

